# Leveraging CryoEM and AI-Driven Morphological Feature Analysis for Insights on Bacterial Structures

**DOI:** 10.64898/2025.12.06.692658

**Authors:** Sita Sirisha Madugula, Lynnicia N. Massenburg, Spenser R. Brown, Amber N. Bible, Chanda R. Harris, Lance X. Zhang, Kiara Parker, Scott T. Retterer, Jennifer L. Morrell-Falvey, Rama K. Vasudevan, Alexis N. Williams

**Affiliations:** Center for Nanophase Materials Sciences, Oak Ridge National Laboratory, Oak Ridge, Tennessee, USA; Biosciences Division, Oak Ridge National Laboratory, Oak Ridge, Tennessee, USA

**Keywords:** Low-dose cryoEM, AI-feature identification, YOLO, microbiology

## Abstract

Bacteria adapt by undergoing dynamic structural changes in response to environmental cues, which are often indicative of deeper phenotypic shifts in physiology and behavior. Understanding these changes across length scales is crucial for elucidating bacterial lifecycles, informing antifouling surface design, enhancing pathogen detection, and advancing renewable energy applications. Cryogenic electron microscopy (cryoEM) enables high-resolution imaging of bacterial structures in hydrated, biologically relevant conditions. However, current quantitative analysis strategies that extract structural information from bacterial samples remain labor-intensive. This work presents an AI-driven segmentation workflow tailored to low-dose cryoEM datasets to rapidly analyze bacterial ultrastructural features from *Pantoea* sp. YR343, a Gram-negative bacterium isolated from the rhizosphere of *Populus deltoides* that forms robust biofilms along plant roots. YOLOv11 image segmentation quantifies inner and outer membrane thickness and flagella length, enabling automated analysis of bacterial ultrastructure. The workflow reliably distinguishes membranes from carbon edges of the TEM grid and contaminant crystalline ice while matching manual measurements and increasing throughput. Flagella detection routines additionally quantify nearest-neighbor proximity between bacterial envelopes and flagella. A field-of-view module further detects bacteria at low magnification for rapid screening. Together, these tools provide a scalable and automated framework for high-throughput, quantitative analysis of bacterial ultrastructure in cryoEM data.

AI-based tools enable rapid characterization of bacterial ultrastructure in low-dose cryoEM. The envelope thickness tool quantifies membrane thickness and anisotropy. The flagella module analyzes filament morphology and detects cell-flagella contacts. The field-of-view (FOV) module

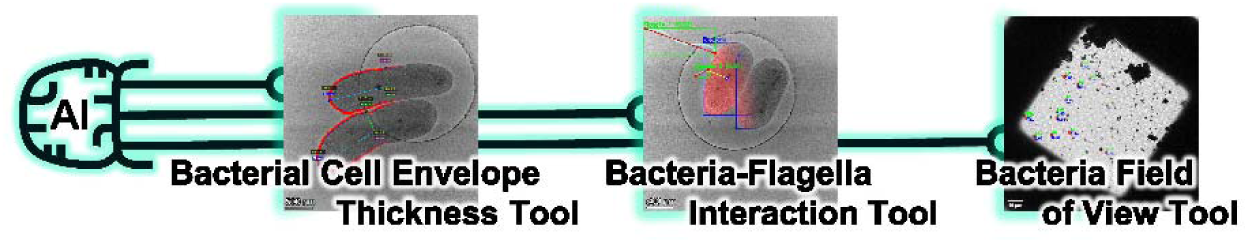

## 1. Introduction

Bacterial morphology is highly diverse and dynamically remodels in response to environmental conditions.^(1)^ These structural variations reflect underlying phenotypic adaptations that influence microbial survival, surface interactions, and function. Understanding these changes is essential for designing antifouling surfaces, improving pathogen detection strategies, and advancing microbial bioenergy applications.^(2)^ Various imaging modalities probe bacterial structures across length scales. Light microscopy (LM) permits live-cell imaging but is limited to ~200 nm resolution. Super-resolution techniques improve this but rely on fluorescent dyes or genetically encoded reporters.^(3)^ Electron microscopy (EM) achieves nanometer resolution but requires fixed, dehydrated, and stained samples that distort native structure. Cryogenic EM (cryoEM) images vitrified samples in near-native, hydrated conditions at high resolution but yields low contrast micrographs with poor signal-to-noise, complicating bacterial segmentation.^(4)^ Traditional edge-detection (e.g. Canny and Sobel) and thresholding methods struggle to delineate cell boundaries in such noisy, heterogenous data.^(5, 6)^

Several segmentation tools exist for LM datasets, combining image correction, edge detection, and morphological annotations.^(7–10)^ These approaches depend on well-defined edges, strong intensity gradients, and user-defined parameters. As a result, they perform poorly on low-dose cryoEM images where edges are faint and intensity gradients are weak. Deep-learning segmentation methods overcome these limitations, handling noisy, low-contrast data beyond the capabilities of traditional algorithms.^(3, 11–14)^ Recent work shows neural networks can recognize subtle bacterial features even under low signal conditions^(13, 15)^ and have improved detection of contaminants in cryoEM images.^(16)^ Vision foundation models such as Meta’s Segment Anything Model (SAM) further demonstrate generalizable segmentation of eukaryotic and prokaryotic cells across imaging modalities.^(14)^ However, to date, no segmentation methodologies have been reported that leverage advanced deep-learning models for quantitative feature extraction directly from 2D cryo-EM micrographs of bacteria. Here, we address this gap by applying the YOLO class of object detection and segmentation models to enable rapid, quantitative bacterial ultrastructure analysis from 2D cryo-EM data.

This work introduces a YOLOv11-based segmentation workflow^(17)^ tailored for low-dose cryoEM bacterial images (**Figure 1**). By leveraging YOLOv11’s high-accuracy instance segmentation capability and curated annotations via Roboflow,^(18, 19)^ the workflow automatically identifies bacteria in noisy micrographs and extracts key morphological features (e.g. cell envelope thickness, flagella, and cell shape). Within this workflow we incorporated three dedicated tools for (1) envelope thickness quantification, (2) analysis of flagella-bacteria overlaps, and (3) bacteria detection within the field of view (FOV) using the Gram-negative *Pantoea* sp. YR343. These tools offer new and innovative approaches to analyze bacterial ultrastructures in cryoEM images. The envelope thickness tool utilizes a 10,000 radial point-pair measurement per cell, enabling fine-grain analysis of cell envelope anisotropy. Benchmarking against manual annotations demonstrates high segmentation accuracy (precision=0.914, recall=1.0, and mAP50=0.995) while delivering a 1,000-fold increase in throughput compared to manual measurements. Training set analysis shows models trained on 50% of the dataset reach stable performance, with minimal improvement beyond the full set, providing practical guidance for deep learning on low-dose cryoEM data. The flagella-interaction and large-field detection tools further accelerate quantification of bacterial structural features. The flagella module measures flagella length and proximity to neighboring bacteria, enabling down-selection of potentially attached flagella for manual verification. The FOV module identifies multiple bacteria at low magnification to rapidly screen surface coverage and identify areas of interest for higher resolution analysis. Together, these tools streamline image analysis, reduce user bias, and enable quantitative morphology studies while improving throughput and reproducibility in low signal to noise cryoEM workflows.

**Figure 1.**
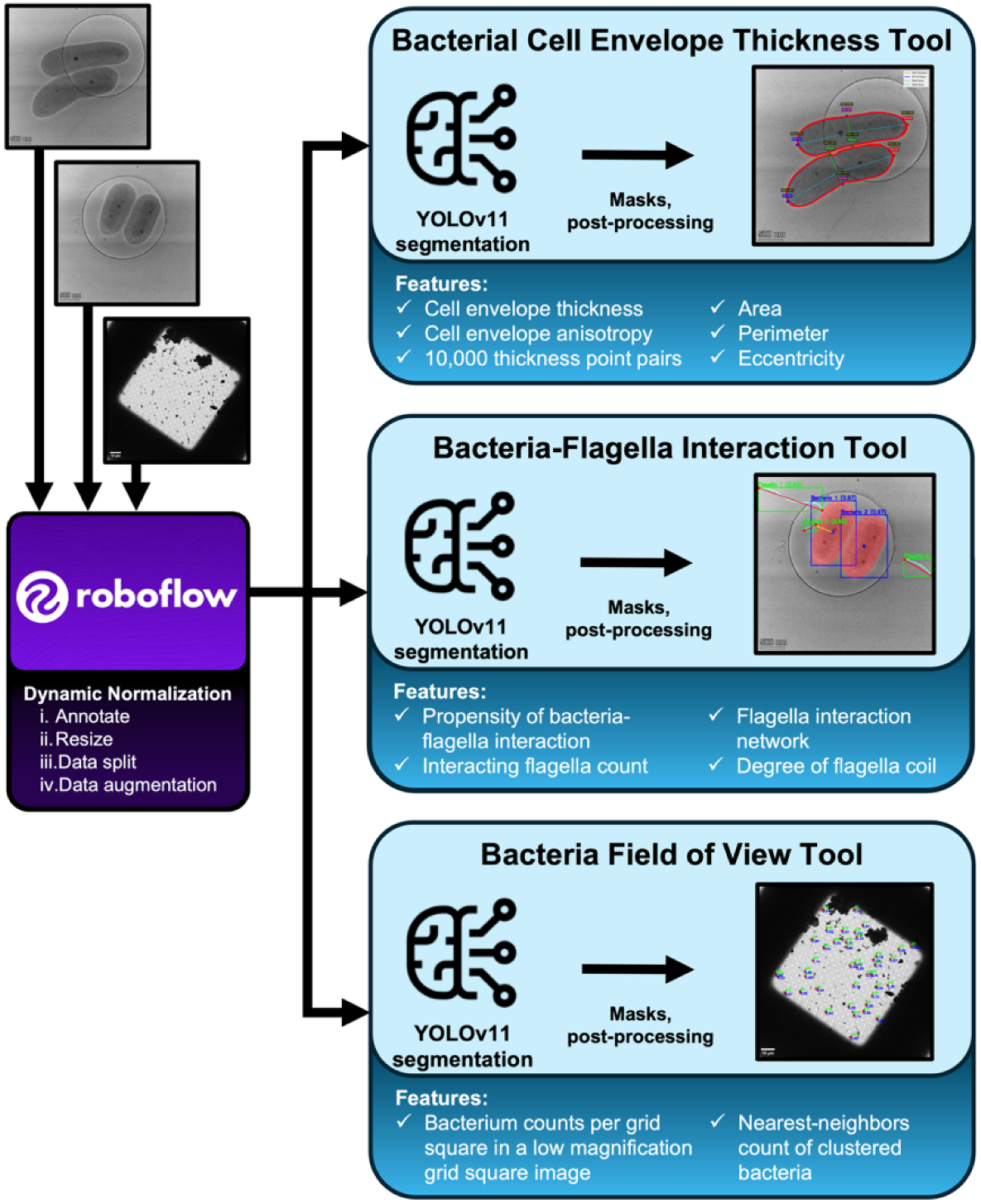
Overview of YOLOv11-based segmentation tools. Low- and high-magnification cryoEM micrographs are annotated and augmented in Roboflow to train YOLOv11 models for three tasks: (1) envelope-thickness and anisotropy from inner/outer-membrane masks, (2) flagella detection with interaction mapping and coiling metrics, and (3) field-of-view (FOV) bacterial detection and counting.

## 2. Results and Discussion

### 2.1 Bacterial Cell Envelope Thickness Tool

We trained YOLOv11 segmentation models to quantify bacterial membrane features, including envelope thickness, anisotropy, area, perimeter, and eccentricity. Precise segmentation enabled direct measurements of envelope thickness, defined as the distance between the outer (OM) and inner (IM) membranes, a key indicator of structural integrity. Details of the training, hyperparameters, and augmentation for model tuning are provided in Section 4. We treated OM and IM contours as distinct classes within a single YOLO model, allowing simultaneous prediction of both membranes with high accuracy.

To implement these capabilities, we developed a workflow that integrates the YOLO-based segmentation model with downstream computational analysis (**Figure 2**). We pre-processed raw cryoEM micrographs of *Pantoea sp.* YR343 grown under different nutrient conditions and used those micrographs to train YOLOv11 models. The trained models predicted OM and IM masks, from which we uniformly sampled 10,000 OM-IM distances per bacterium and then converted those values into nanometer-scale measurements (**Figure S1**). The addition of pole and side annotations along the major cell axis enabled analysis of anisotropy and correlations between envelope thickness and shape descriptors, laying the foundation for the quantitative comparisons.

**Figure 2.**
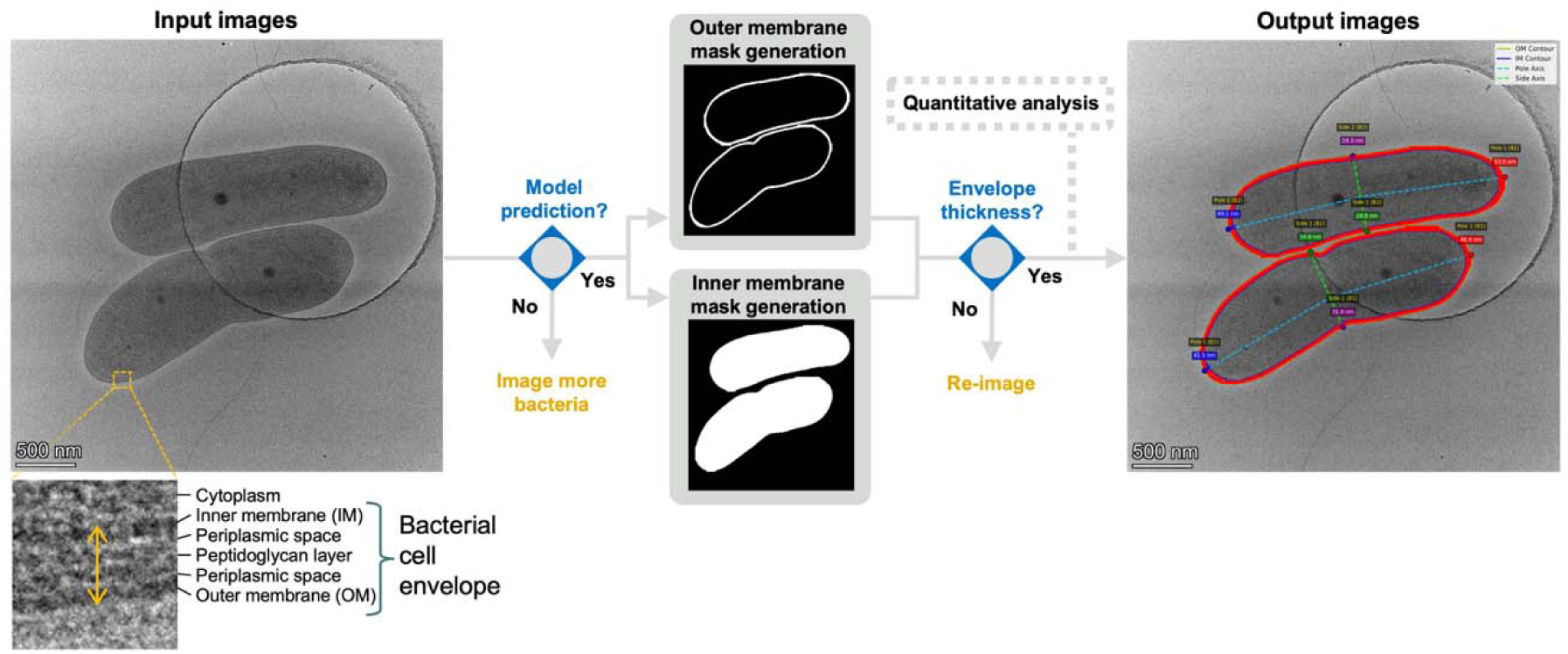
Envelope-thickness workflow and output. Low-dose cryoEM images of *Pantoea* sp. YR343 are segmented with YOLOv11 to generate outer- and inner-membrane masks. For each cell, the tool samples 10,000 OM-IM point pairs to compute thickness (nm) and annotates poles (cyan) and sides (green) for anisotropy analysis. Scale bar: 500 nm.

We evaluated model performance on validation and test sets and observed high predictive accuracy for both the OM and IM classes. Precision and recall were closely matched across the two classes, indicating reliable identification of membrane regions with minimal false detections (**Table S1** and **Table S2**). The mAP50 values and F1 scores showed strong alignment between predicted and annotated masks, demonstrating that our model maintained robustness on the low signal-to-noise micrographs in identifying the fine structures of bacterial membranes. The precision-recall and F1 curves for the validation sets are shown in **Figure S2**.

#### 2.1.1. Biological Relevance of Bacterial Cell Envelope Thickness

We applied the envelope thickness tool to determine how various growth media altered bacterial morphology. The tool quantified thickness, anisotropy, area, perimeter, and eccentricity across growth conditions. Bacteria grown in minimal MOPS (3-(N-morpholino)propanesulfonic acid) media supplemented with either glucose (preferred carbon source that promotes glycolysis) or succinate (non-preferred carbon source that promotes gluconeogenesis) display similar morphologies when compared to bacteria grown in the nutrient rich R2A media. To control for growth phase, all cultures were imaged at the stationary phase using cryoEM and LM (**Figure S3** and **S4**). Both imaging modalities showed compact, round cells in R2A and elongated rods in MOPS supplemented with either carbon source. The cryoEM data revealed the MOPS grown culture exhibited thicker envelopes and larger areas than the R2A samples. Quantitative analysis confirmed these trends (**Table 1**). Cells grown in R2A were smaller and rounder (e.g., displayed lower eccentricity and smaller perimeter) than those in MOPS. Eccentricity correlated positively with cell area and perimeter (**Figure S5**), showing that cell enlargement was driven primarily by elongation. Mean envelope thickness only showed a weak positive correlation with eccentricity, indicating that cell elongation outpaced envelope thickening under nutrient-limited conditions. These analyses reflect representative cryoEM datasets acquired under each growth condition, with multiple individual bacteria measured per medium types (n of 37-51). These datasets were used to demonstrate the workflow’s quantitative precision rather than serve as biological replicate-to-replicate variability.

**Table 1.** Morphological parameters of *Pantoea* sp. YR353 under low-dose cryoEM conditions during stationary phase in R2A, MOPS + glucose, and MOPS + succinate media.

| Media (bacteria count) | Cell envelope thickness (nm)*<sup>□</sup> | Area (μm<sup>2</sup>)* | Perimeter (μm)* | Eccentricity* |
| --- | --- | --- | --- | --- |
| R2A (n=51) | 28.33 ± 3.41 | 0.92 ± 0.16 | 4.27 ± 0.41 | 0.46 ± 0.17 |
| MOPS + glucose (n=37) | 46.34 ± 10.30 | 1.76 ± 0.45 | 6.91 ± 1.90 | 0.89 ± 0.06 |
| MOPS + succinate (n=50) | 25.90 ± 8.68 | 1.21 ± 0.26 | 5.49 ± 0.96 | 0.86 ± 0.06 |

\*Values represent the mean ± standard deviation.
<sup>□</sup> A total of 10,000 cell envelope thickness distances were measured per whole imaged bacteria.

Radial analysis revealed clear anisotropy between poles and sides (**Figure 3a**). Both MOPS grown cultures showed thicker poles, with the effect strongest in the glucose supplemented samples. This is consistent with the higher mean thickness reported in **Table 1**. Violin plots of thickness distribution (**Figure 3b**) showed broader upper tails for the minimal media cultures, reflecting increased variability among anisotropic cells.

**Figure 3.**
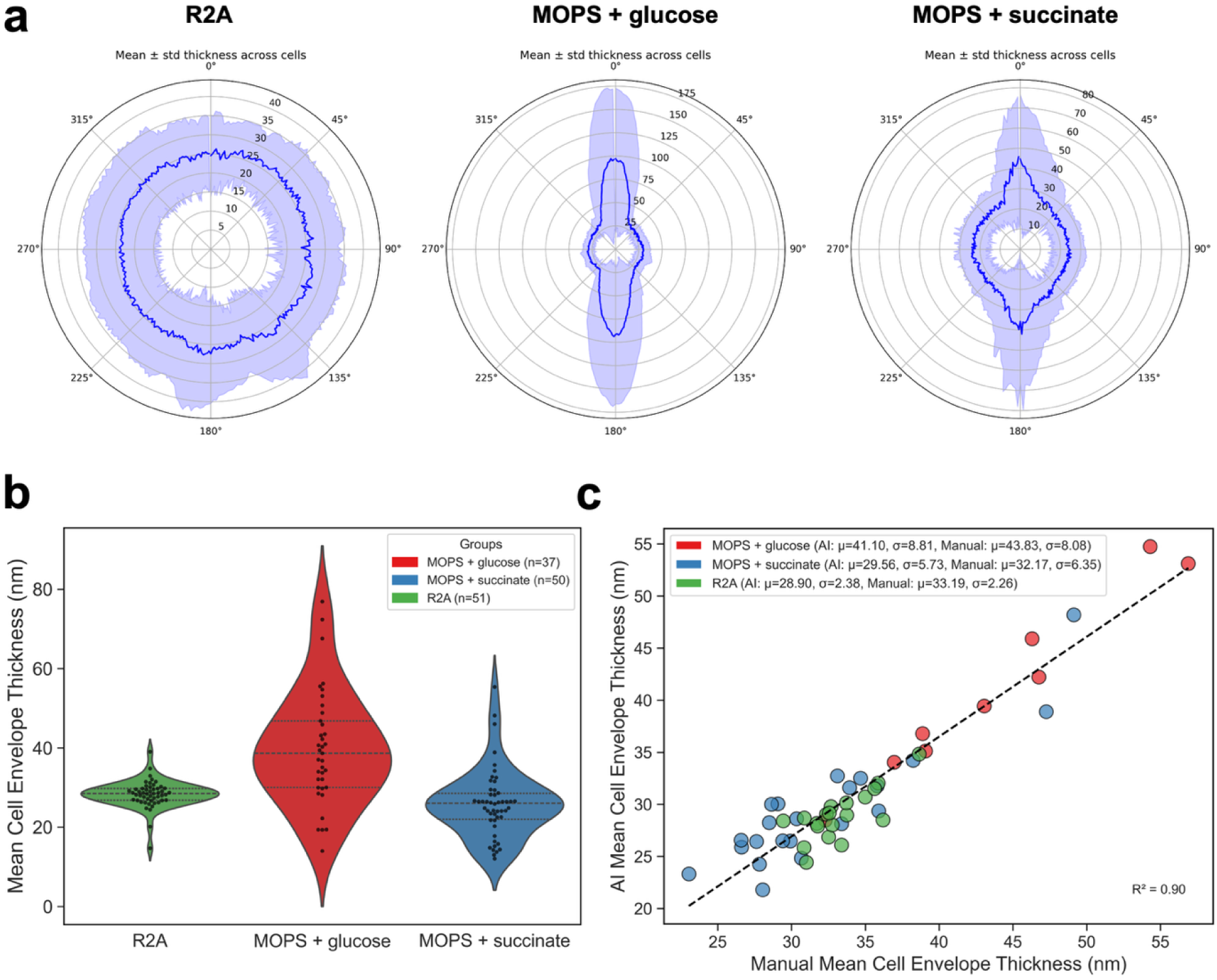
Quantitative analysis using the bacterial cell envelope thickness tool. (**a**) Radial plots of *Pantoea* sp. YR343 envelope thickness in R2A, MOPS + glucose, and MOPS + succinate media. The 0° position marks pole 1, followed counterclockwise by side 1 (90°), pole 2 (180°), and side 2 (270°). (**b**) Violin plots showing global thickness distributions across media types. (**c**) Correlation between AI- and manual-based thickness measurements (n=50).

These observations indicate that, in our dataset, *Pantoea* sp. YR343 altered its envelope architecture under different nutrient conditions. Cells grown in rich R2A media appear more spherical, while those grown in minimal media, independent of carbon source, retain the traditional rod-like morphology. However, anisotropy and localized thickening can vary in the minimal media dependent on carbon source. These observations in shape and envelope variability suggest that both parameters are responsive to nutrient availability. This morphological flexibility may extend to other bacterial species, including pathogens, where changes in cell shape could influence surface adhesion and interface interactions.^(20–23)^ Further studies with biological replicate datasets and complementary biochemical assays must occur to confirm these trends and clarify their physiological basis.^(24–28)^ This heterogeneity in bacterial ultrastructure may translate to other bacterial species and could influence surface adhesion, biofilm formation, and overall environmental adaptability.^(29, 30)^

#### 2.1.2. Comparison between AI and Manual Cell Envelope Thickness Measurements

We validated our AI-derived segmentation approach against manual segmentation (n=50). The two methods correlated strongly with R^2^=0.90 (**Figure 3c** and **S6**). Minor discrepancies occurred near overlapping cells or image artifacts. However, the model correctly distinguishes membranes from carbon hole edges and crystalline ice contaminants (**Figure S1-S6**). **Figure S1a** and **Figure S1b** show instances where cell envelopes contacted the carbon hole edge, yet the model properly detected the lower contrast membrane and the higher contrast carbon hole edge. **Figure S6b** shows crystalline ice contamination does not affect the analysis. **Figure 3c** illustrates the high correlation between the model and manual measurements. These results highlight the strength of this tool in speed, detection accuracy, and reliable performance on low-dose cryoEM images.^(31–33)^

### 2.2. Bacteria-Flagella Interaction Tool

We further deployed YOLOv11 segmentation models on low-dose bacterial micrographs to detect flagella. **Figure 4** shows the outline for flagella detection method with example outputs shown in **Figure S7**. The workflow used similar features of interest as the envelope thickness tool to identify bacteria outer membranes with an additional feature set to detect flagella. This tool required the bacterial OM model as the initial reference. When trained with only flagella annotations, segmentation was limited to images lacking bacteria. The model successfully identified free, non-interacting flagella as well as interacting flagella, achieving precision=0.947, recall=1.0, mAP50=0.995, and mAP50-95=0.746 (**Figure S8**, **Table S1** and **Table S3**). Given the resolution ranges and focus of this workflow, we define interacting flagella as instances where flagella overlap with or appear close to bacteria membrane. Interacting flagella are flagged for manual review, allowing the end user to perform further analysis to determine if flagella are attached. **Figure S7** shows the model accurately detected flagella of varying lengths and contrast levels, independent of bacterial morphology. This robust performance underscored the effectiveness of automated annotation and segmentation in reducing the labor-intensive nature of manual identification for low contrast features of interest.

**Figure 4.**
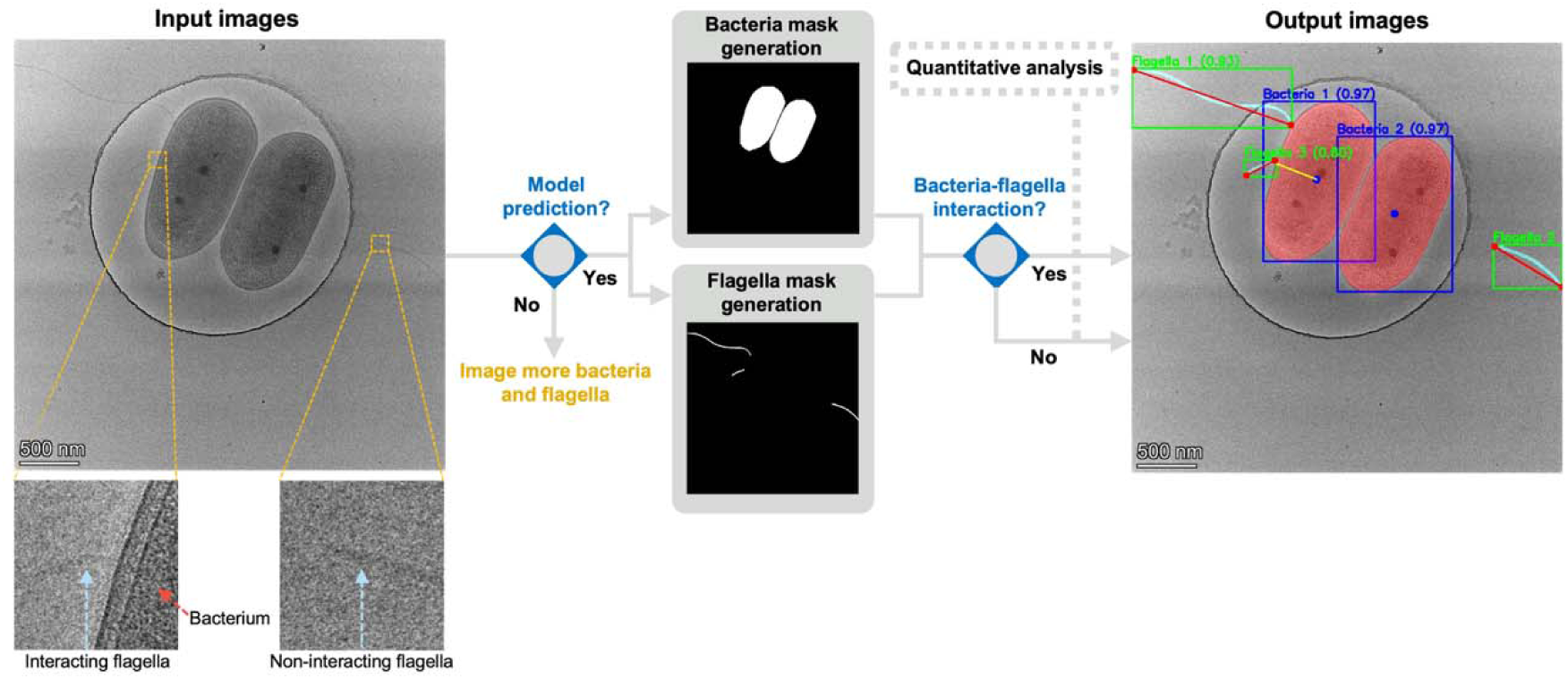
Bacteria-flagella overlap workflow and output. Low-dose cryoEM images of *Pantoea* sp. YR343 are segmented with YOLOv11 to generate outer-membrane and flagella masks. Insets show interacting versus non-interacting filaments. Overlap occur when the flagellar mask intersects the bacterial OM mask. Output overlays display bacteria (blue boxes; red OM masks) and flagella (green boxes; cyan masks). Scale bar: 500 nm.

#### 2.2.1. Biological Relevance of Flagella Interactions

Next, we used the bacteria-flagella interaction tool to find trends in flagella interactions and degree of coiling. Detecting flagella interactions is particularly valuable for high-throughput screenings to evaluate bacterial motility and surface adhesion. The tool rapidly quantified the number of interacting flagella per bacterium, constructed a global flagella interaction network, and detected flagella morphology (**Figure 5a,b** and **Figure S9a,b**). Global trends in the current dataset illustrated the type of population-level analyses enabled by this workflow. Most flagella were free, non-interacting filaments on the carbon film, with only a small fraction overlapping with bacterial envelopes (**Figure 5a,b**). This observation is possibly due to partial flagella detachment, weak flagella attachment, or limited imaging area. Among interacting cases, most bacteria exhibited fewer than five associated flagella, though a minority displayed markedly higher interaction frequency. Network analysis revealed individual flagella could contact multiple neighboring bacteria, demonstrating the workflow can resolve higher-order connections and local clustering behavior. *Pantoea* sp. YR343 typically carries a single polar flagellum, with occasional secondary polar filaments.^(21)^ Our tool captures these sparse, low-contrast flagella interactions and can assist in identifying potential rare cross-cells associations in the data that would be difficult to detect manually. Such interactions may underlie early stages of surface adhesion or biofilm formation and will be useful in future biological studies.

**Figure 5.**
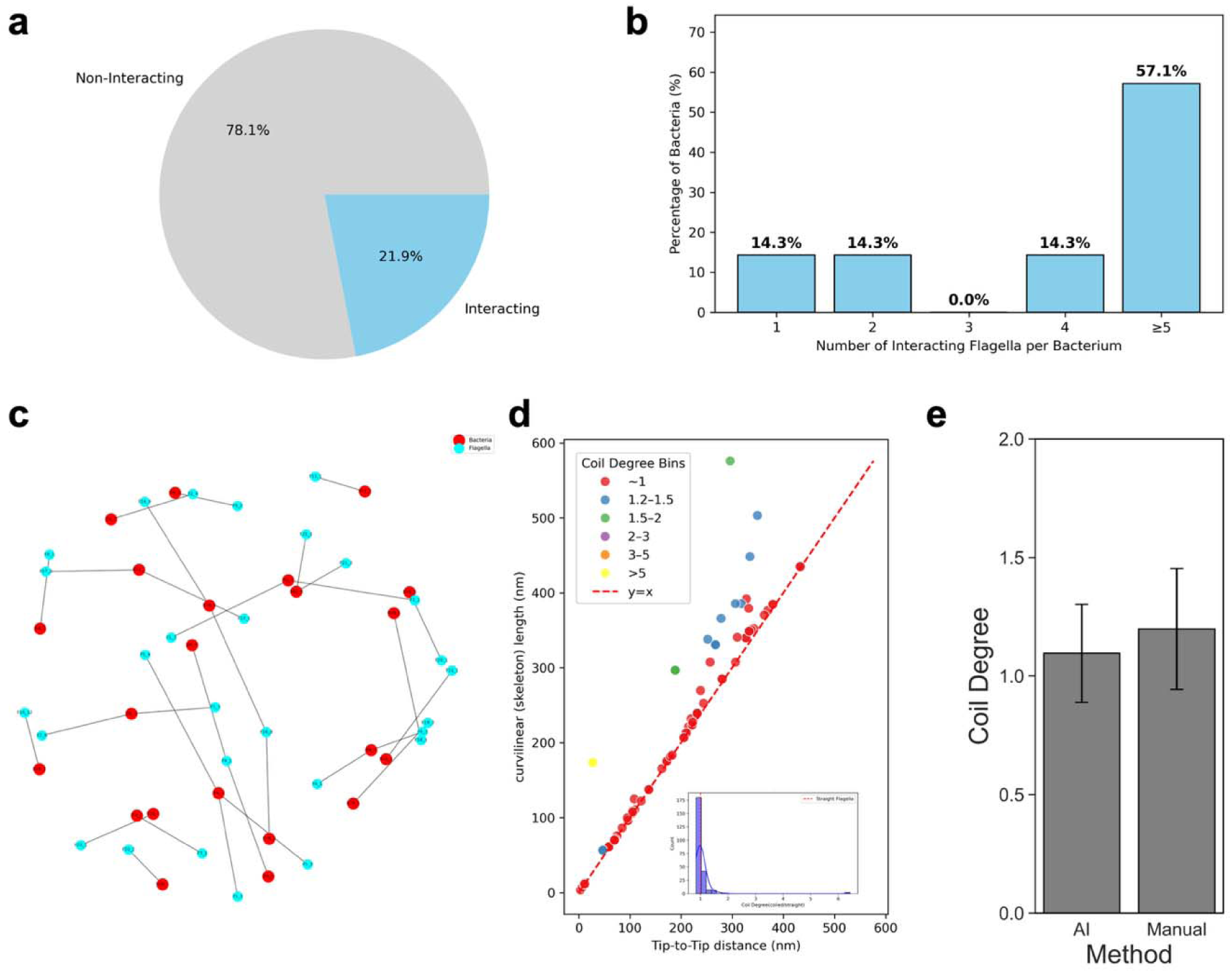
Quantitative outputs from the bacteria-flagella overlap tool. (**a**) Proportion of flagella that interact with bacteria. (**b**) Distribution of interacting flagella per bacterium. (**c**) Degree of coiling for individual flagella with inset histogram of coiling frequency. (**d**) Mean coiling degree measured by AI and manual tracing (n=10).

Beyond interaction mapping, the tool also quantified flagellar morphology. Here we introduced a metric termed the “degree of coiling”, defined as the ratio of flagellar length to the tip-to-tip distance (**Figure 5c** and **Figure S9c**).^(34, 35)^ Ratios near 1 corresponded to uncoiled, extended filaments, whereas values ≥ 5 indicated strongly coiled flagella. Our results revealed most flagella are predominantly uncoiled, with only a few exhibiting significant coiling (**Figure 5c**). Both long and short flagella occasionally displayed high coiling, but such cases were rare. Variations in apparent flagella length is likely due to partial truncation within the image frame. Importantly, the ratio-based metric compensates for truncation effects introduced by the image frame, allowing coiling trends to be assessed independently of flagellar length and comparisons across images and magnifications. This measure facilitates high-throughput screening of surface interactions that alter flagellar adhesion and coiling behavior, processes linked to early biofilm formation.^(36–40)^ Together, these analyses demonstrate our automated pipeline provides rapid, scalable, and reproducible population-level insights into flagellar organization, interaction networks, and structural states. These capabilities are challenging to achieve through manual analysis alone.^(41–43)^

#### 2.2.2. Comparison between AI and Manual Attached Flagella Length Measurements

The AI model accurately measured flagella length and tip-to-tip distance to determine the degree of coiling. We compared the AI degree of coiling to the manual measurements (n=10) and found that the two methods agree (**Figure 5e**). Both methods are effective at observing the uncoiled, elongated nature of flagella observed in the cryoEM images (**Figure S10**). Moreover, the AI-based approach provided the added advantage of simultaneously capturing the bacteria perimeter and area, enabling integrated correlation analysis between cell morphology and flagellar architecture. These results demonstrate that automated flagella quantification achieves accuracy comparable to manual tracing while providing higher throughput and improved reproducibility.

### 2.3. Bacteria Field of View Tool

A third component of our workflow identified multiple bacteria within a single grid-square FOV (**Figure 6** and **Figure S11**). This model reliably detected individual cells under low-dose conditions, even when contaminating crystalline ice or debris were present. Performance metrics confirmed high accuracy and confidence scores (> 0.75) across micrographs (**Figure S12**, **Table S1** and **Table S4**). This ability to function under low-dose conditions was particularly significant as it allows large-scale, unbiased surveys of microbial populations even when image quality was compromised by dose limitations. Automating this step minimizes manual inspection and enables high-throughput morphological screening at the population level.

**Figure 6.**
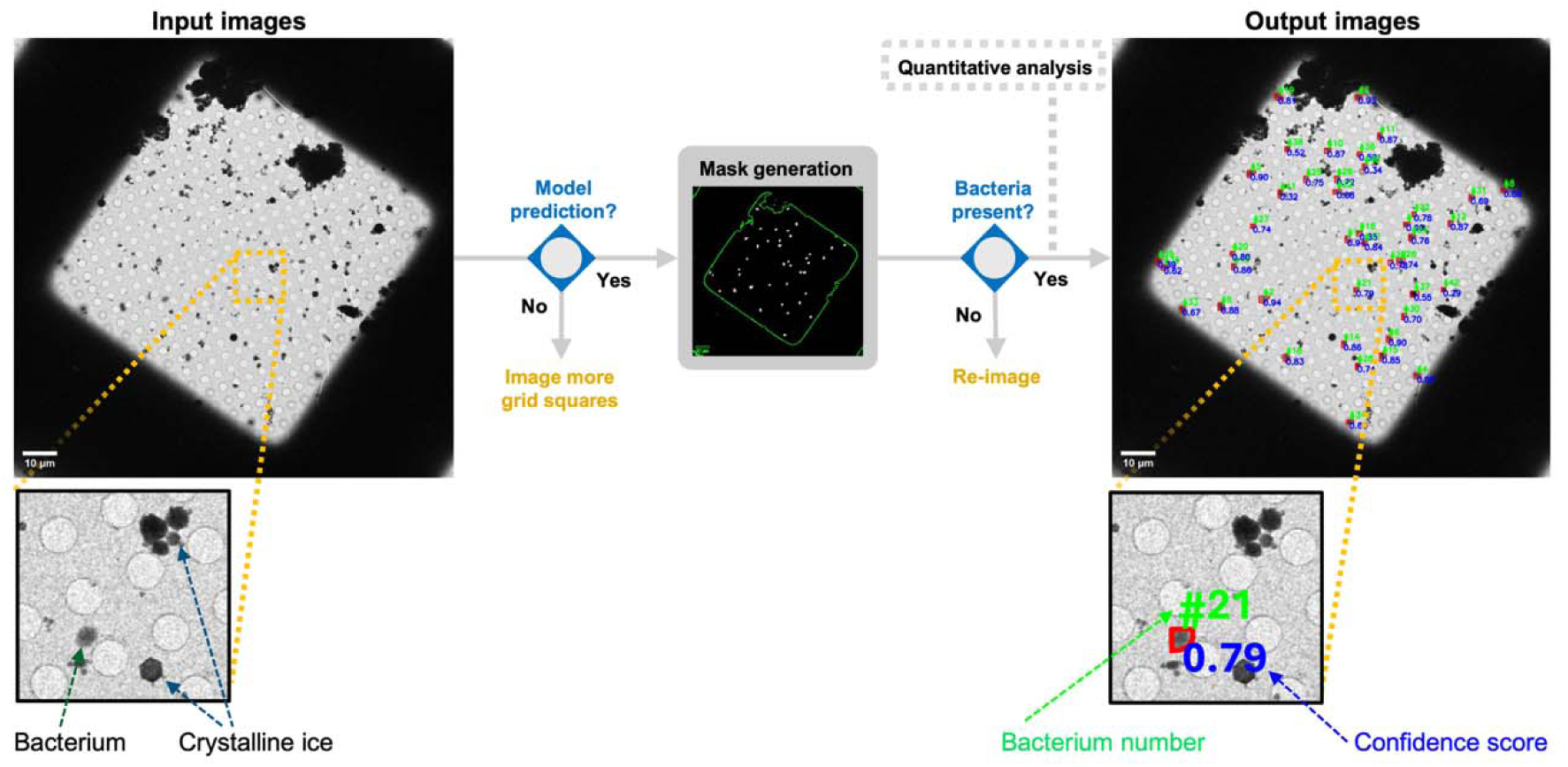
FOV workflow and output. Low-magnification cryoEM grid-square images of *Pantoea* sp. YR343 are segmented with YOLOv11 to generate bacterial masks in the presence of ice crystals. Output overlays display per-cell IDs (green) and confidence scores (blue). Insets highlight bacteria versus ice crystals in the larger FOV images. Scale bar: 10 µm.

#### 2.3.1 Data Screening Relevance of Bacteria in a Field of View

This bacteria FOV tool provides a quantitative “birds-eye-view” of the bacterial sample quality for each grid square imaged at 260 × magnification. Lower magnification grid square images have lower resolution, and the bacteria “blobs” may be mistaken for ice crystals (**Figure 7a**). To address possible misidentification, search map montages consisting of stitched images of higher magnification (3.8 k×) data, were used to verify bacterial presence (**Figure 7b**). The identified bacteria were then annotated in the corresponding lower magnification grid square images in the pre-processing steps to ensure training occurred only on bacteria. We chose the lower 260 × magnification for grid square imaging in this workflow to optimize for speed of data screening and allow for future integration with other automated cryoEM software (e.g., EPU and SerialEM). The tool effectively counted bacteria in various configurations, including single cells and clusters, across different experimental conditions (**Figure 7c** and **Figure S11**). A nearest-neighbor analysis quantified bacterial clustering within 1 µm of each centroid, showing greater clustering in MOPS + succinate media compared to MOPS + glucose and R2A (**Figure S13**). This clustering difference was likely due to sample preparation differences in blotting time and not necessarily correlated to media type as the bacterial ODs were similar during cryoEM preparation. This analysis provides a rapid means of evaluating grid quality and optimizing imaging targets. Low magnification grid square images were the ideal for this tool because each image could be captured in only a few seconds, compared with more than 15 minutes required to collect a search map montage of the same area. Therefore, the bacteria FOV tool provides rapid grid square counting and grid-square level nearest-neighbor population analysis to enhance the cryoEM screening process.^(44–47)^

**Figure 7.**
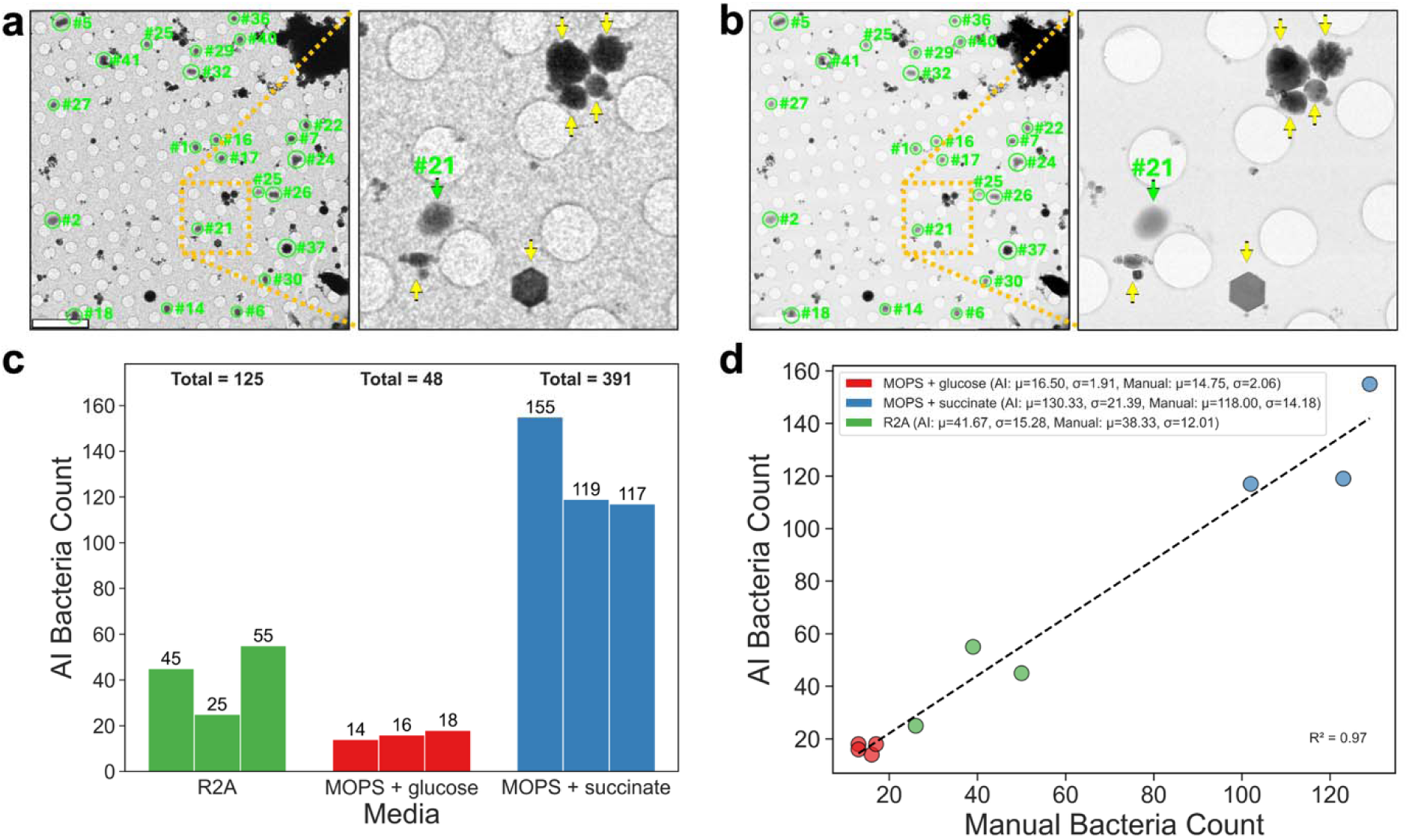
Quantitative analysis using the FOV tool. (**a**) Low-magnification grid-square cryoEM image of *Pantoea* sp. YR343 in R2A at 260 × showing bacteria (green circles/arrows) and ice crystals (yellow arrows) in vitreous ice. Scale bar: 10 µm. (**b**) Search-map montage at 3.8 k× showing higher-resolution views of bacteria and cubic/hexagonal ice. (**c**) AI-derived bacterial counts per grid square across media. (**d**) Correlation between AI- and manual-based counts across grid squares (R²=0.97).

#### 2.3.2 Comparison between AI and Manual Counts of Bacteria in a Field of View

To check the accuracy of the bacteria FOV tool, the AI-based counts were compared to the manual counts of bacteria on a 260 × magnification grid square (n=10) (**Figure 7d**). The two methods correlated strongly (R^2^ of 0.97), confirming accuracy and reproducibility (**Figure S14**). Overall, the fast-imaging workflow does not compromise the bacteria-picking accuracy of the FOV tool, underscoring its reliability even in low signal-to-noise grid square images.

## 3. Conclusion

This AI-based workflow demonstrates substantial potential for broad applications in microbiology, bio-interface research, and healthcare.^(48–51)^ It enables rapid bacterial characterization, reduces manual intervention, and accelerates ultrastructure analysis. YOLOv11-based models mitigate artifact segmentation from carbon edges and crystalline ice contamination and perform effectively across diverse object sizes, from flagella to entire cells. These tools may be extended to other Gram-negative bacteria, such as *E. coli*, with minimal retraining for species-specific features. Application to Gram-positive bacteria will require model retraining to account for differences in cell wall morphology.^(51, 52)^ However, the model’s ability to distinguish elongated and spherical morphologies (Section 2.1) suggests that it could also recognize coccoid Gram-positive species such as *S. aureus* with minimal retraining and validation. Future development will focus on integrating these models into automated cryo-EM acquisition pipelines using SerialEM to enable on-the-fly bacterial segmentation and feedback during data collection. This integration would streamline high-throughput screening workflows, allowing real-time detection of structural phenotypes and adaptive targeting of regions of interest for downstream analysis.

## 4. Materials and Methods

### Bacterial Growth

*Pantoea* sp. YR343 wild-type (WT) is a Gram-negative bacterium initially isolated from the rhizosphere of native *Populus deltoides* trees in North Carolina.^(21)^ Bacterial strains were cultured on R2A agar (BD Biosciences) at 29 °C. Single colonies were used to inoculate liquid cultures in either rich R2A broth (R2A Broth Premix, TEKnova, Inc.), minimal MOPS medium supplemented with 0.4% glucose (MOPS + glucose) or minimal MOPS medium supplemented with 0.4 succinate (MOPS + succinate). Cells were harvested and analyzed at stationary phase (24 hr growth) for each culture.

### Confocal Microscopy

*Pantoea* sp. YR343 cells were grown to stationary phase in R2A, MOPS + glucose and MOPS + succinate media. For each condition, 20 µL cells were imaged on glass microscope slides using a 100 × objective on a Zeiss 710 Confocal Scanning Laser Microscope.

### CryoEM Sample Preparation

For stationary growth conditions, samples were collected from R2A, min+glucose, or min+succinate cultures after 24 hr growth. For each condition, 3 µL aliquots were applied to glow-discharged Quantifoil® or C-Flat 2/2 gold grids, blotted using a VitroBot Mark IV (Thermo Fisher Scientific, blot force −5 for 5-10 s at 100% humidity and 4.5 °C), and vitrified by plunge-freezing in liquid ethane/propane cooled by liquid nitrogen.

### CryoEM Data Collection

Manual image collection utilized standard low-dose (40 e^−^/Å², −5 μm defocus) with a 70 μm C2 aperture at cryogenic temperatures on a Thermo Fisher Scientific Krios G4 with a Falcon 3EC direct electron detector (Thermo Fisher Scientific, counting mode). Magnifications ranged from 260 × (30.960 nm/pix microprobe TEM mode) used to capture bacteria at a field of view grid square to 8.7 k× (0.899 nm/pix, nanoprobe TEM mode) used to capture bacterial cell envelope thickness and flagella length with EPU (Thermo Fisher Scientific) and Velox (Thermo Fisher Scientific). Search maps (7 × 7 image montages) were collected at 3.8 k× (2.128 nm/pix nanoprobe TEM mode) with Tomo5 (Thermo Fisher Scientific) to confirm the presence of bacteria or crystalline ice contamination on the grid square images collected at 260 × magnification.

### Manual Measurements

Manual measurements were conducted using 3dmod (v4.11.25) for cryoEM images. A total of five manual measurements were times for each tool to benchmark the manual throughput speed compared to the AI throughput speed. For all manual measurements, the scale bar was also manually measured with one open contour with two points to track the pixel size of each image. To compare the measurements of the bacterial cell envelope tool, the OM to IM distance was manually measured as one open contour with two points. A total of 10 points were selected radially around the bacteria perimeter in 36° increments from 0° counterclockwise. AI pole 1 position at 0° and centroid position were manually aligned onto the cryoEM image to create a similar starting position for the manual 10-point pair measurement comparison. A radial plot tool was created to guide the manual placement of 10 points in 36° increments (the manual measurements were as follows: 0°, 36°, 72°, 108°, 144°, 180°, 216°, 252°, 288°, 324°) to keep the angle placement independent of AI (**Figure S6**). To compare measurements of the bacteria-flagella interaction tool, flagella that were proximal to the bacteria are considered as interacting with the bacteria regardless of flagella attachment state. Interacting flagella counts were isolated as points with one contour per bacteria. Interacting flagella lengths were measured as the perimeter of a series of points tracking the flagella filament. The tip-to-tip measurement was measured as two points from the start and end of the flagella (the flagella at the edge of an image frame was considered the end point of the flagella even if the flagella is likely to extend beyond the image frame). The ratio of flagella coiling was determined by dividing the length distance by the tip-to-tip distance. To compare the bacteria counts in the bacteria FOV tool, individual points were placed on each bacterium on a 260 × magnification grid square and counted per image. The 3.8 k× search maps were used as a guide for bacteria identification on the grid square (**Figure 7a,b**).

### AI-based Measurement Method for OM-IM Membrane Thickness

IM and OM masks were generated by a segmentation model and processed to calculate periplasmic thickness. For each bacterium, OM and IM contours were extracted and the centroid of the IM was determined. Principal Component Analysis (PCA) of the OM contour is used to identify the long axis (poles) and perpendicular short axis (sides) of the bacterial cell. Uniform sampling all along the OM contour was then performed, and for each sampled OM point, the shortest Euclidean distance to the IM contour is calculated using a k-d tree nearest-neighbor search. These distances represent local IM-OM thickness measurements and were annotated with the corresponding angular orientation relative to the pole axis. This yielded a complete angular thickness profile around the bacterium. In addition, values at predefined angular intervals (0°, 36°, …, 324°) as well as pole and side positions were extracted for reference. Distances are then converted to nanometers using the imaging scale, and results were saved as per-cell CSV files containing both the full set of 10,000 measurements and the angle-specific summaries, alongside annotated images showing contours, centroid, axes, and OM-IM rays.

### AI-based Measurement Method for Calculating Degree of Coil

To obtain the degree of coil of the flagella, the flagellar masks obtained from the flagella detection tool are skeletonized to reduce each filament to a one-pixel-wide medial axis. The curved length of each flagellum is measured by summing skeleton pixels and converting to nanometers using the imaging scale. The two terminal endpoints of each skeleton are identified as tips, and their Euclidean distance is measured to obtain the uncoiled tip-to-tip length. The degree of coil is then calculated as the ratio of the curved skeleton length to the direct tip-to-tip distance, with values close to 1 indicating nearly uncoiled flagella and higher values reflecting increased coiling.

### Preprocessing and Data Annotation

Raw bacterial cryoEM 4096 × 4096 images are adaptively contrast-normalized using variance-based percentile scaling and saved as lossless 8-bit PNGs to preserve fine cryoEM feature. Thereafter, these images are manually annotated using the Roboflow platform.^(19)^ Three complementary annotation tasks are performed to capture distinct ultrastructural features. For the cell envelope thickness model, the OM and IM of each bacterium are annotated as polygon contours. For flagella detection, polygon annotations are generated for both bacterial outer membranes and flagella filaments, enabling analysis of flagella-cell association. For the multiple bacteria in FOV detection model, polygon annotations are drawn around all bacteria present in each FOV image, allowing automated quantification of cell density and spatial organization. Datasets are exported in YOLOv11 format (TXT polygon/box annotations with accompanying YAML configuration files) for direct use in the Ultralytics training pipeline. Task-specific resizing, augmentations and pre-processing summarized in **Table 2**, are applied to account for variability limited image availability. To maximize training diversity given the limited number of expert-curated images, the flagella and FOV dataset are split into 80% training, 10% validation, and 10% test sets. To maximize training diversity given the limited number of expert-curated images, the flagella and FOV dataset are split into 80% training, 10% validation and 10% test sets, while the thickness model has a 80:5:15 ratio (80% training, 5% validation and 15% test).

**Table 2.** Summary of datasets, augmentations, and resizing applied for model building.

| S. No | Model | Initial Data Size | Augmented Data Size | Augmentations and Resizing |
| --- | --- | --- | --- | --- |
| 1 | Cell envelope thickness | 149 | 403 | Horizontal and vertical flip, crop 0% minimum and 20% maximum zoom, rotation between $-15^{\circ}$ and $+15^{\circ}$, $\pm 10^{\circ}$ horizontal, $\pm 10^{\circ}$ vertical shear, saturation between $-25\%$ and $+25\%$, resized to $640 \times 640$ |
| 2 | Flagella identification | 275 | 713 | Horizontal and vertical flip, crop-0% minimum zoom, 15% maximum zoom, rotation between $-15^{\circ}$ and $+15^{\circ}$, $\pm 10^{\circ}$ horizontal and $\pm 10^{\circ}$ vertical shear, saturation between $-25\%$ and $+25\%$, brightness between $-15\%$ and $+15\%$ resized to $1024 \times 1024$ |
| 3 | Bacteria detection in a FOV | 50 | 120 | Horizontal and vertical flip, Crop-0% minimum zoom, 20% maximum zoom, $\pm 10^{\circ}$ horizontal and $\pm 10^{\circ}$ vertical shear, saturation between $-25\%$ and $+25\%$, blur up to $2.5 \times$, resized to $1024 \times 1024$ |

### YOLOv11 Model Training and Hyperparameters

YOLOv11n-seg (image segmentation, IS) models are fine-tuned in the Ultralytics framework using Google Colab Pro with GPU acceleration (PyTorch backend) for all the three main prediction tasks discussed in the study. Pretrained YOLOv11n-seg weights are used for initialization and models are trained with task-specific hyperparameters. The thickness and flagella segmentation models are trained for 300 epochs with a batch size of 16, learning rate of 0.001, and IoU threshold of 0.5, while the bacteria FOV model was trained for 200 epochs with the same learning rate. All models employed the AdamW optimizer with a cosine learning rate schedule (final lrf=0.01). Early stopping is applied with a patience of 20 epochs for the flagella model and 50 epochs for the FOV model, whereas the thickness model was run for the full 300 epochs until convergence. Model validation and performance are monitored using loss functions, precision-recall curves (PR curves), F1-scores and mean average precision (mAP). The best-performing weights are retained for downstream analyses.

### Model Evaluation

Model performance was quantitatively assessed using multiple metrics, including mean Average Precision at IoU thresholds of 0.5 (mAP50), precision, recall, and F1-score. Training progress and model convergence are monitored using several loss functions: box loss for evaluating localization accuracy, segmentation loss for pixel-level accuracy, classification loss for accurate object identification, and distribution focal loss (DFL) to improve bounding box predictions. These comprehensive evaluations ensured reliable and robust model performance. Model training data for instance segmentation and evaluation can be found in **Table 2**.

## Acknowledgements

The authors gratefully acknowledge Brian Ofsthus for his assistance with cryoEM database management and seamless data transfer from the microscope facility to the data center. This work is supported by the U.S. Department of Energy, Office of Science FWP ERKCZ64, Structure Guided Design of Materials to Optimize the Abiotic-Biotic Material Interface, as part of the Biopreparedness Research Virtual Environment (BRaVE) initiative. A portion of L.M.’s effort was supported by the Center for Nanophase Materials Sciences (CNMS), which is a U.S. Department of Energy, Office of Science User Facility at Oak Ridge National Laboratory. Microscopy was performed using instrumentation within ORNL’s Materials Characterization Core provided by UT-Battelle, LLC, under Contract No. DE-AC05-00OR22725 with the DOE and sponsored by the Laboratory Directed Research and Development Program of Oak Ridge National Laboratory, managed by UT-Battelle, LLC, for the U.S. Department of Energy.

## Data Availability Statement

All datasets are openly available on Constellation (DOI: 10.13139/ORNLNCCS/2997581).

## Code Availability

The models and all analysis/training scripts are available at [GitHub: https://github.com/Sireesiru/Cryo-EM-Ultrastructures].

