## Supplemental Data for "Leveraging CryoEM and AI-Driven Morphological Feature Analysis for Insights on Bacterial Structures"

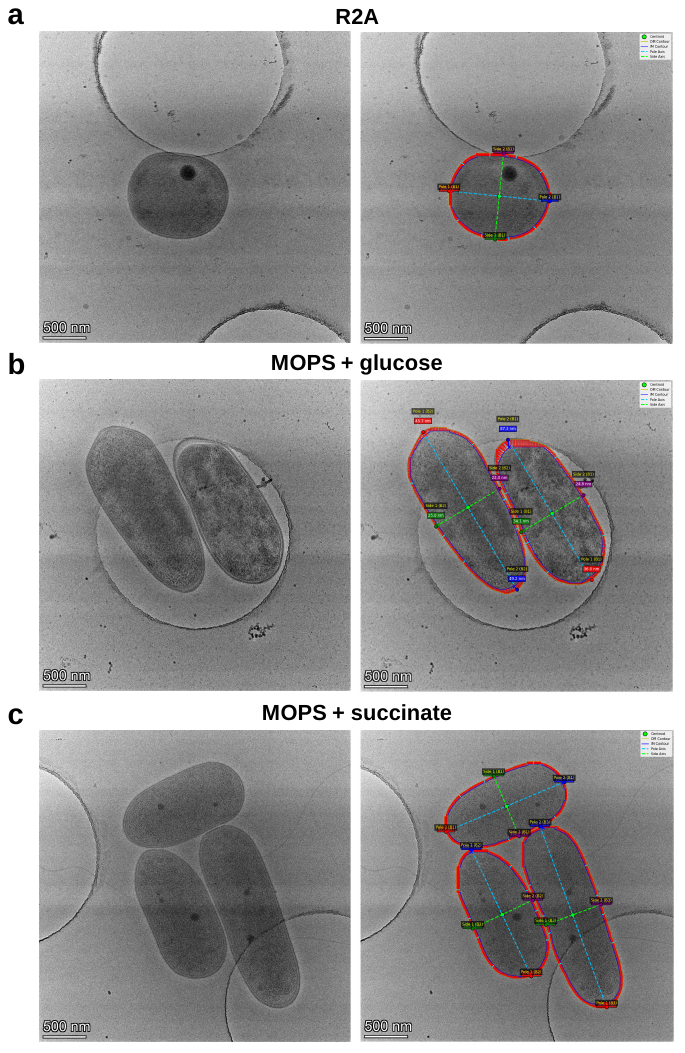

**Figure S1.** Example outputs of bacteria structure analysis with the bacteria cell envelope thickness tool. Input (left) and output (right) cryoEM images of *Pantoea* sp. YR343 grown in (**a**) R2A, (**b**) MOPS + glucose and (**c**) MOPS + succinate. The tool is compatible with a single bacterium, bacteria pairs and bacteria clusters. Red lines represent the 10,000 point pairs around the cell envelope perimeter. Green dashed lines represent the bacteria sides and cyan dashed lines represent the bacteria poles. Scale bar is 500 nm.

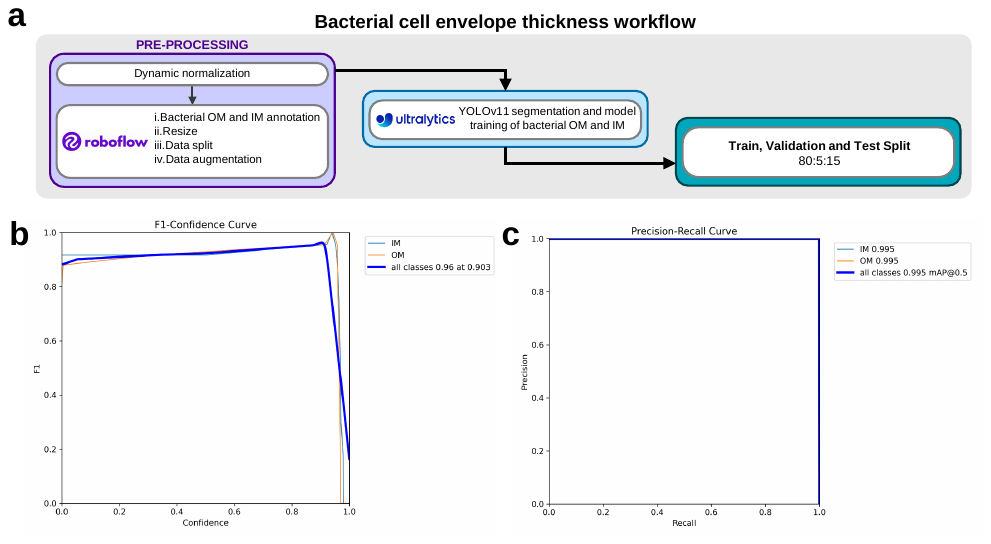

**Figure S2.** Validation curves for the bacterial segmentation model used in the bacteria cell envelope thickness tool. (**a**) Bacteria-flagella interaction workflow with Roboflow preprocessing and YOLOv11 model training, validation and testing. (**b**) Precision-Recall curve and (**c**) F1-Confidence plot for the bacterial envelope thickness detection model.

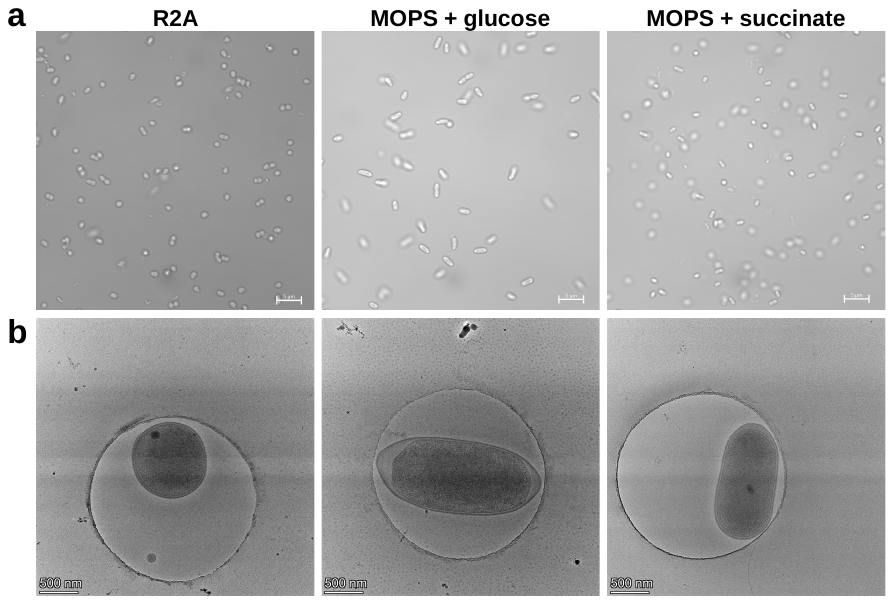

**Figure S3.** Morphology of *Pantoea* sp. YR343 grown in different media. (**a**) Confocal microscopy images (scale bar is 5 µm) and (**b**) cryoEM images (scale bar is 500 nm) of *Pantoea* sp. YR343 grown in R2A, MOPS + glucose and MOPS + succinate.

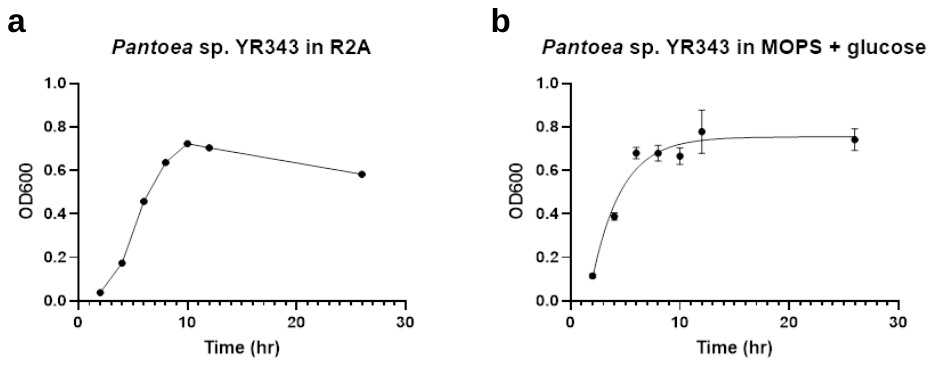

**Figure S4.** Growth curves of *Pantoea* sp. YR343 grown in different media. (**a**) *Pantoea* sp. YR343 grown in R2A and (**b**) MOPS + glucose.

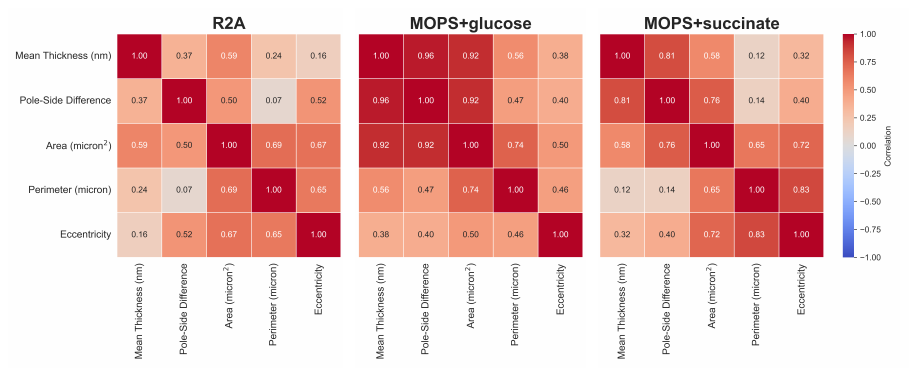

**Figure S5.** Correlation heat maps of bacterial features of interest. Correlation heat maps of *Pantoea* sp. YR343 bacterial cell features based on media of R2A, MOPS + glucose or MOPS + succinate. Correlation heat maps show the positive or negative correlation between bacteria bacterial cell envelope mean thickness, pole-side difference, area, perimeter and eccentricity of bacteria.

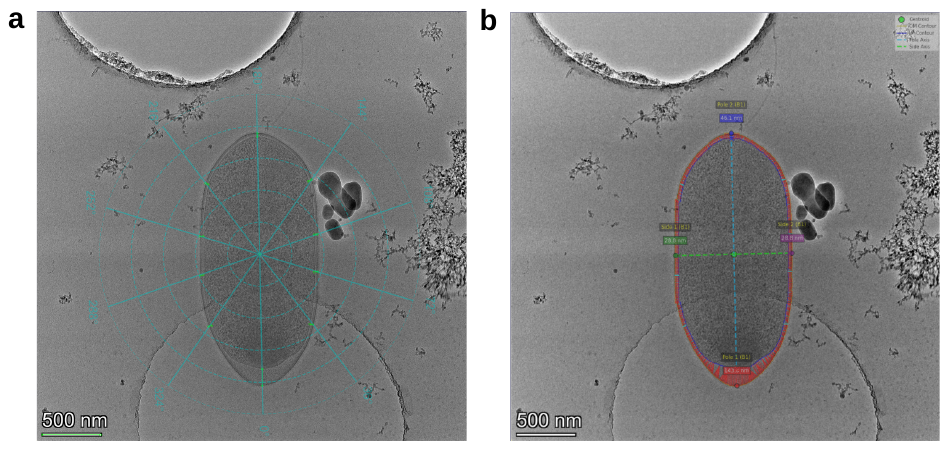

**Figure S6.** Measurements of bacterial membrane thickness. (**a**) Radial plot ruler used to chart angles for manual cell envelope measurement. The following angles (cyan lines) were used for the manual measurements: 0°, 36°, 72°, 108°, 144°, 180°, 216°, 252°, 288°, 324°. The intersecting points between the radial blue lines and the outer membrane counterclockwise (CCW) from pole 1 (0°) were used as the cell envelope thickness markers to compare to (**b**) AI-based measurements. Scale bar is 500 nm.

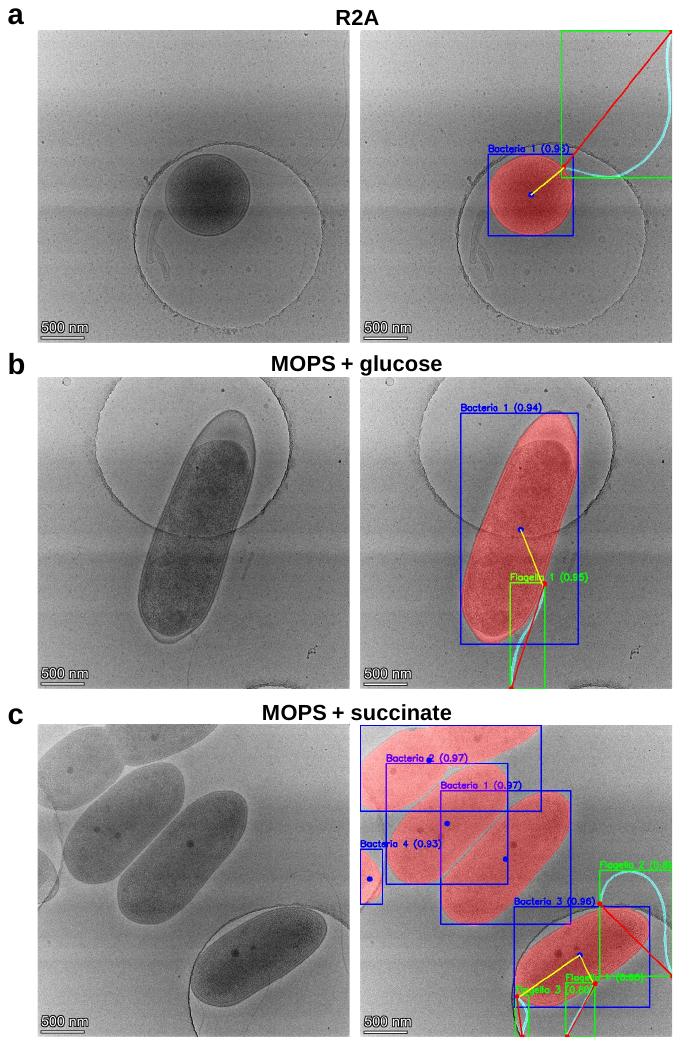

**Figure S7.** Example outputs of bacteria structure analysis with the bacteria-flagella interaction tool. Input (left) and output (right) cryoEM images of *Pantoea* sp. YR343 grown in (**a**) R2A, (**b**) MOPS + glucose and (**c**) MOPS + succinate. The tool is compatible with a single bacterium, bacteria pairs and bacteria clusters. Bacteria segmentation is highlighted in red with a blue boundary box with the confidence score. Flagella are segmented in cyan with a light green boundary box and confidence score. The yellow lines represent the location of interaction between flagella per bacterium and red lines represent the tip-to-tip distance from the bacteria-flagella interaction point to the end of the flagella (or the flagella ends for a non-interacting flagellum shown in **Figure 4**). Scale bar is 500 nm.

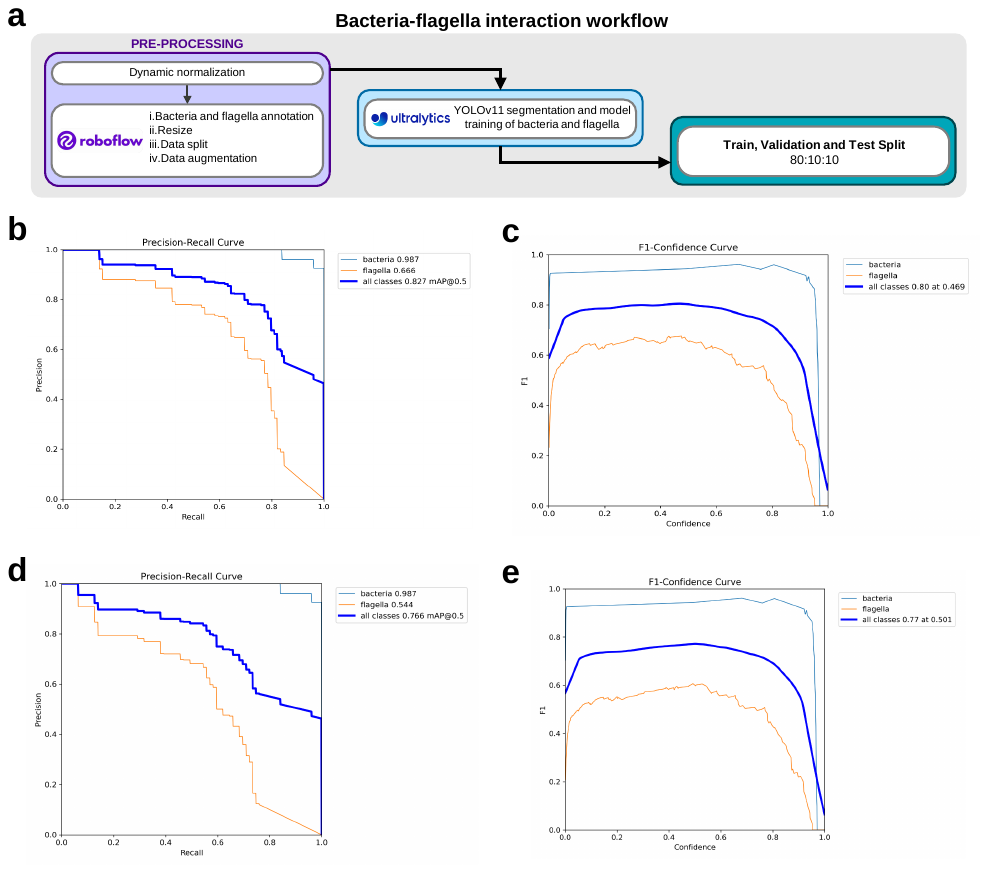

**Figure S8.** Validation curves for the bacteria and flagella segmentation models within the bacteria-flagella overlap tool. (**a**) Bacteria-flagella overlap workflow with Roboflow preprocessing and YOLOv11 model training, validation and testing. (**b**) Box Precision-Recall (**c**) Box F1-Confidence (**d**) Mask Precision-Recall and (**e**) Mask F1-Confidence plots for the bacteria-flagella overlap segmentation model.

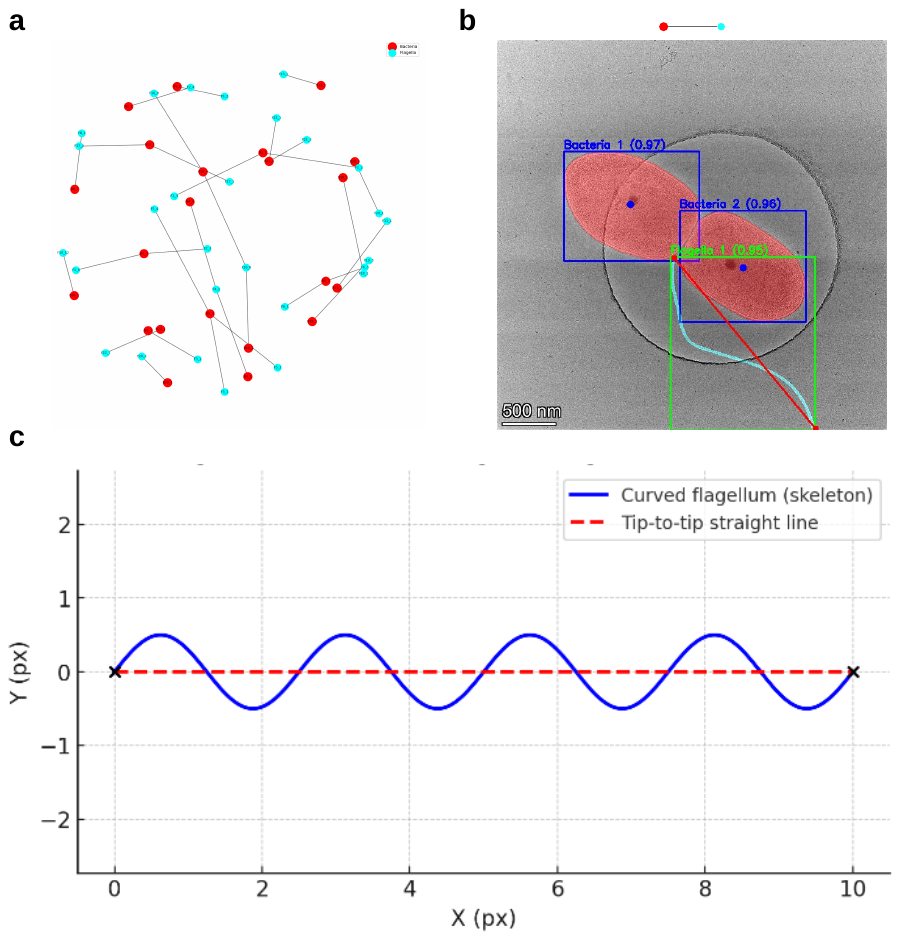

**Figure S9.** Bacteria-flagella interaction map and flagella degree of coil quantification. (**a**) Bacteria-flagella interaction map showing connectivity of flagella among neighboring bacteria across 73 8.7kx images. (**b**) An example image showing one bacteria-flagella interaction and example illustrated interaction map above the image. (**c**) Schema of flagellum degree of coil calculation. Curvilinear (blue) vs. tip-to-tip (red) lengths. Tips are represented as a black cross on either ends of the blue curve.

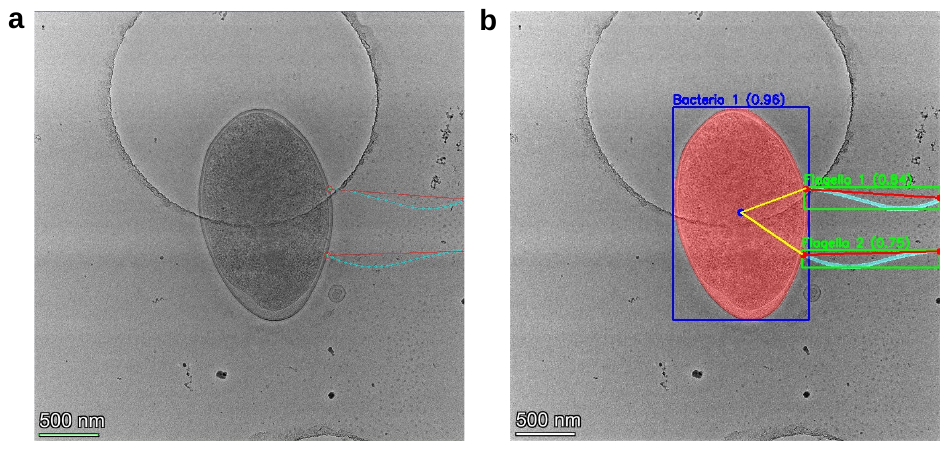

**Figure S10.** Measurements of interacting flagella lengths and tip-to-tip distances. (**a**) Manual flagella lengths (cyan contour lines) measured with the tip-to-tip distances (red contour lines) to compare to (**b**) AI-based measurements. Scale bar is 500 nm.

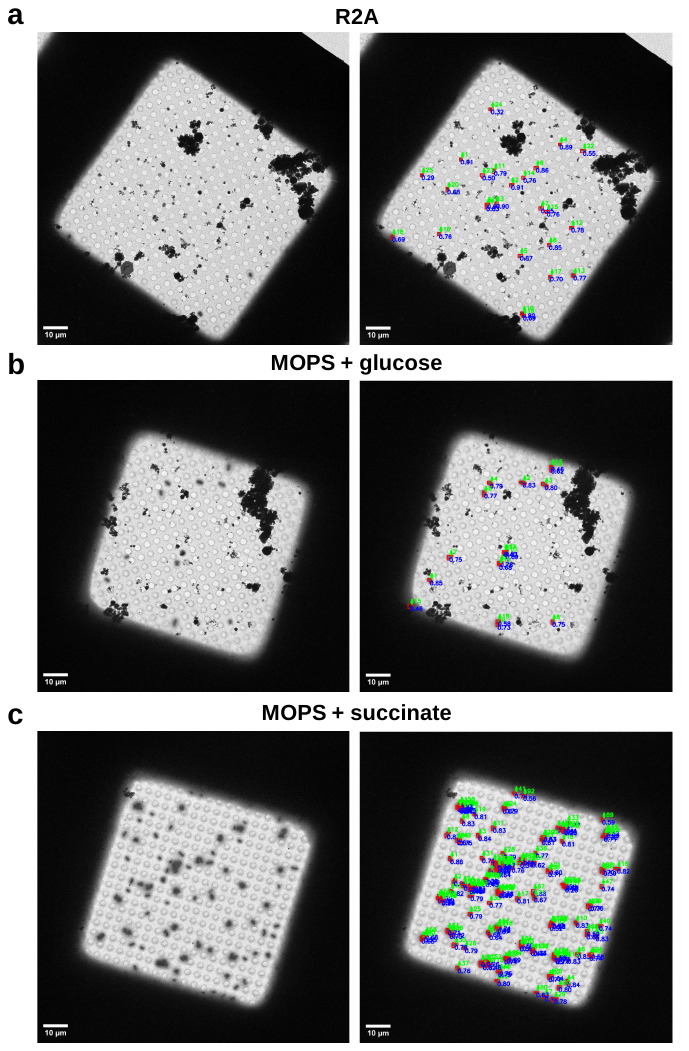

**Figure S11.** Example outputs of bacteria structure analysis with the bacteria FOV tool. Input (left) and output (right) cryoEM images of *Pantoea* sp. YR343 grown in (**a**) R2A, (**b**) MOPS + glucose and (**c**) MOPS + succinate. Bacteria segmentations are shown in red tracing the bacterial perimeter. The bacterial count (light green text) and confidence score (blue text) are also output for each bacterium segmentation. The tool is compatible with a single bacterium, bacteria pairs and bacteria clusters. Scale bar is 10 µm.

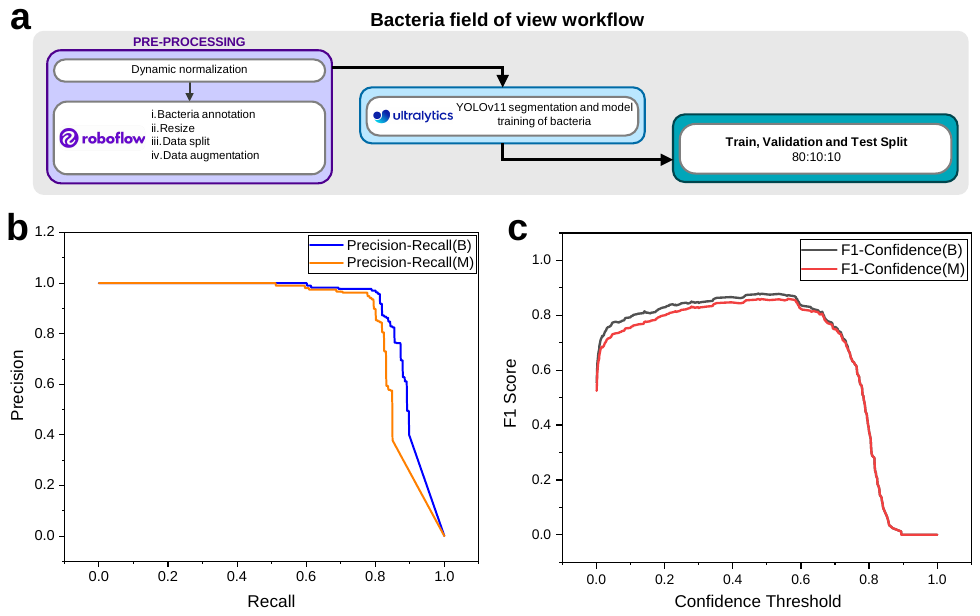

**Figure S12.** Workflow and validation curves for the bacteria FOV tool. (**a**) Bacteria field of view workflow with Roboflow preprocessing and YOLOv11 model training, validation and testing. (**b**) Precision-Recall curves and (**c**) F1-Confidence plots for the bacteria FOV detection model.

**Figure S13.** Nearest neighbor distribution plot by media type. (**a**) Nearest neighbor distribution plot of the bacteria distribution by media type in 10 grid squares (k=4). A closer neighbor distance represents a higher clustering density of the bacteria at the centroid x,y coordinates for each bacterium. Example nearest neighbor distances are shown as grey lines connecting between bacteria marked with (**b**) cyan points on the MOPS + succinate medium grid square, (**c**) light green points on the R2A medium grid square and (**d**) red points on the MOPS + glucose grid square. The grid squares are arranged from highest (left) to lowest (right) bacteria clustering density. The original grid square images are shown in **Figure S11**.

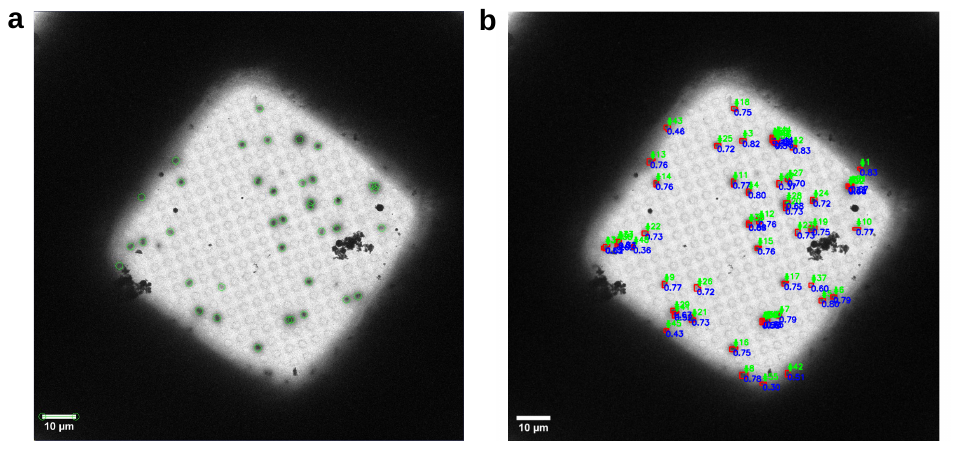

**Figure S14.** Measurements of bacteria identification. (**a**) Manual bacteria counts as individual contour points to compare to (**b**) AI-based bacteria counts. Scale bar is 500 nm.

**Table S1.** Performance of the three models on their respective test sets.

| **Model** | **Precision** | **Recall** | **F1** | **mAP@0.5** |
| --- | --- | --- | --- | --- |
| Thickness model | 0.952 | 1 | 0.995 | 0.995 |
| Flagella model | 0.896 | 0.837 | 0.886 | 0.878 |
| FOV model | 0.693 | 0.605 | 0.632 | 0.611 |

**Table S2.** Performance metrics for the YOLOv11 Low-dose thickness model on the validation set.

| **S. No.** | **Mask Precision** | **Mask Recall** | **mAP50** |
| --- | --- | --- | --- |
| Overall | 0.914 | 1.0 | 0.995 |
| Inner Membrane | 0.913 | 1.0 | 0.955 |
| Outer membrane | 0.915 | 1.0 | 0.995 |

**Table S3.** Performance metrics for the YOLOv11 bacteria-flagella interaction segmentation model.

| **S. No.** | **Box**  **Precision** | **Box**  **Recall** | **mAP50**  **(Box)** | **Mask Precision** | **Mask**  **Recall** | **mAP50 (Mask)** |
| --- | --- | --- | --- | --- | --- | --- |
| Overall | 0.807 | 0.816 | 0.827 | 0.778 | 0.774 | 0.766 |
| Bacteria | 0.891 | 1.000 | 0.987 | 0.894 | 1.000 | 0.987 |
| Flagella | 0.723 | 0.633 | 0.666 | 0.663 | 0.547 | 0.544 |

**Table S4.**  Performance metrics for the YOLOv11 FOV model.

| **S. No.** | **Box**  **Precision** | **Box**  **Recall** | **mAP50**  **(Box)** | **Mask Precision** | **Mask Recall** | **Mask mAP50** |
| --- | --- | --- | --- | --- | --- | --- |
| Low-dose FOV model | 0.964 | 0.803 | 0.894 | 0.945 | 0.783 | 0.856 |
